# Dopaminergic learning and arousal circuits mediate opposing effects on alcohol consumption in *Drosophila*

**DOI:** 10.1101/624833

**Authors:** Shamsideen A. Ojelade, Andrew R. Butts, Collin B. Merrill, Eve Privman Champaloux, Yoshinori Aso, Danielle Wolin, Roberto U. Cofresi, Rueben A. Gonzales, Gerald M. Rubin, B. Jill Venton, Aylin R. Rodan, Adrian Rothenfluh

**Affiliations:** Department of Psychiatry, Program in Neuroscience, UT Southwestern Medical Center, Dallas, TX; Molecular Medicine Program, University of Utah, Salt Lake City, UT; Department of Chemistry, University of Virginia, Charlottesville, VA; Janelia Research Campus, HHMI, Ashburn, VA; Division of Pharmacology and Toxicology, College of Pharmacy, University of Texas at Austin, TX; Department of Internal Medicine, Division of Nephrology, University of Utah, Salt Lake City, UT; Department of Human Genetics, University of Utah, Salt Lake City, UT; Department of Psychiatry, Department of Neurobiology and Anatomy, University of Utah, Salt Lake City, UT

## Abstract

The response to drugs of abuse is a combination of aversive and reinforcing reactions. While much is known about the role of dopamine in mammalian drug reinforcement, we know little about the brain circuits mediating drug aversion. Here we show that two distinct dopaminergic circuits mediate reinforcing and acute aversive responses to alcohol consumption in *Drosophila*. Protocerebral anterior medial dopamine neurons projecting to the mushroom bodies are required for flies to acquire alcohol preference. Conversely, a bilateral pair of dopamine neurons projecting to the dorsal fan-shaped body (dFSB) mediates acute alcohol avoidance. Alcohol consumption can be reduced by decreasing the activity of the appetitive reinforcement-circuit to the mushroom bodies, or by increasing activity in the dopamine neurons projecting to the dFSB. Thus, distinct dopaminergic pathways can be targeted to reduce the intake of harmful drugs.

## Introduction

Alcohol exposure causes both pleasurable and negative responses in humans. People displaying increased sensitivity to alcohol’s rewarding effects or resistance to the acute aversive responses are at increased risk for alcohol use disorder (AUD)^1^. The development of AUD involves circuits mediating the reinforcing effects of alcohol, including dopaminergic projections from the ventral tegmental area to the nucleus accumbens^2^. However, much less is known about the neurons mediating the acute aversive responses to this drug.

Similar to mammals, *Drosophila* show naïve aversion to alcohol when given a choice between liquid food with or without alcohol^3^. This initial alcohol aversion transforms into experience-dependent preference after an alcohol pre-exposure^3^. Dopaminergic neurons are involved in the recall of ethanol-conditioned odor preference^4^. However, it is not known how dopamine is involved in voluntary ethanol consumption in *Drosophila*—including naïve aversion and the transformation from aversion to experience-dependent alcohol preference. Here we show that distinct dopamine circuits mediate the acute aversive and reinforcing effects upon ethanol exposure. We also show that normal ethanol responses require the *Drosophila* dopamine D1R1 receptor in the respective target neurons of these two dopamine circuits, and that manipulations of either circuit can lead to reduction in voluntary alcohol consumption.

## Results

### Distinct dopamine neurons mediate opposing consummatory reactions to alcohol

To determine the involvement of dopamine in alcohol-consumption behavior, we fed flies the dopamine precursor L-DOPA (L-3,4-dihydroxyphenylalanine), or 3-IY (3-iodo-tyrosine), an inhibitor of the rate-limiting dopamine synthesis enzyme tyrosine hydroxylase (TH, Fig. 1a). Flies with increased dopamine levels (Supplementary Figure 1a) showed enhanced naïve aversion, while dopamine depletion (Supplementary Figure 1a) led to naïve preference in an abbreviated 16-hour “2-bottle choice” CAFÉ assay (for capillary feeder, Fig. 1B^5^). As expected (see Fig. 1a), feeding of the TH product L-DOPA was able to restore the naïve alcohol preference caused by TH inhibition back to aversion (Supplementary Figure 1b). After a 20-min alcohol pre-exposure a day prior to testing, control flies developed experience-dependent preference for alcohol, whereas flies with increased dopamine levels did not (Fig. 1b). These results suggested that dopamine is crucial for alcohol aversion rather than acquired preference in flies. To confirm that levels of dopamine signaling correlated with alcohol aversion, we genetically altered DAN activity using a *TH-Gal4* driver, which is expressed in most DAN (Fig. 1c,^6–9^). Enhancing *TH-Gal4* DAN activity (using the heat-activated TrpA cation channel) during pre-exposure and during the CAFÉ assay preference test (Fig. 1d) caused alcohol aversion regardless of pre-exposure (Fig. 1e). Conversely, using a heat-activated dominant negative dynamin (*shi*^*ts*^) to silence these *TH-Gal4* DAN led to alcohol preference, regardless of pre-exposure (Fig. 1e). Therefore, changing the activity of the majority of DAN using *TH-Gal4* had the same effect as pharmacologically altering dopamine levels (Fig. 1b).

**Fig. 1.**
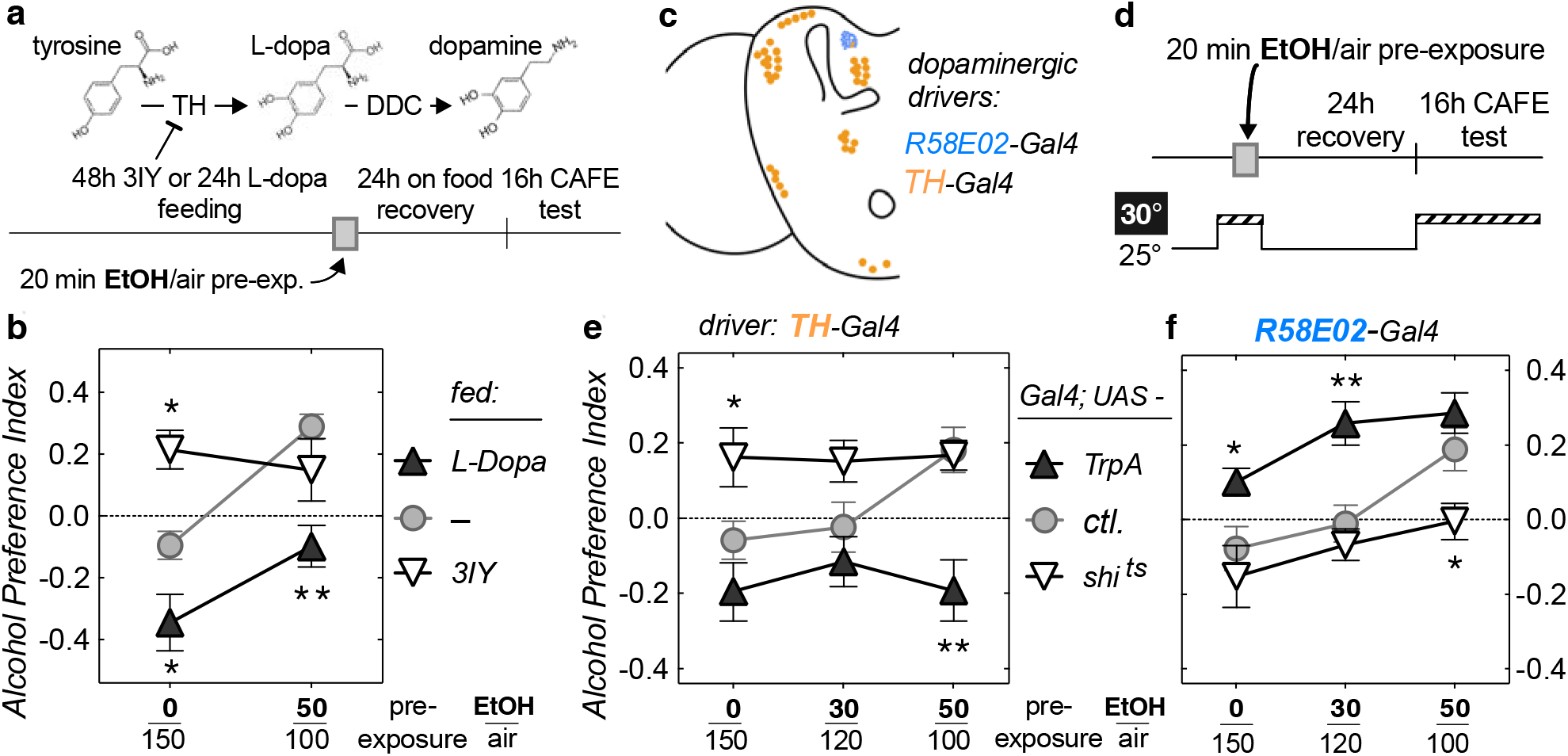
Distinct dopamine neurons are required for naïve avoidance and experience-dependent alcohol preference. (**a**) Experimental paradigm describing drug feeding followed by alcohol-induced consumption preference assay (CAFÉ stands for capillary feeder). (**b**) Naïve control flies (0/150 alcohol/air flow mock pre-exposure, fed plain food: – grey) show slight aversion to alcohol (Preference Index < 0); naïve flies with increased dopamine levels (fed L-DOPA: black) show enhanced aversion; flies with reduced dopamine levels (fed 3IY: white) show naïve preference. Alcohol pre-exposure (to 50/100 alcohol/air flow for 20 min the day before) induces preference in control flies, but not in L-DOPA-fed flies (\*\**P* < 0.01, \**P* < 0.05, two-way ANOVA with Dunnett’s post-hoc comparisons vs. – control). (**c**) Schematic indicating cell bodies of dopamine neurons in the fly brain and the two dopaminergic drivers used in **e**,**f**. (**d**) Experimental paradigm, where 30° C is the effective temperature causing silencing (*shi^ts^*), or activation (*TrpA*) of neurons. (**e**) Silencing *TH-Gal4* dopamine neurons (white) causes naïve preference, and activating them (black) suppresses experience-dependent preference, similar to the pharmacological intervention in **b** (\*\**P* < 0.01, \**P* < 0.05; here, and in following Figures *ctl*. are *UAS-GFP* flies). (**f**) Conversely, activating PAM dopamine neurons (*R58E02-Gal4* driver, black) facilitates experience-dependent preference, while silencing these neurons (white) prevents preference acquisition (\*\**P* < 0.01, \**P* < 0.05).

The PAM cluster (protocerebral anterior medial) of DAN expresses TH, but lacks *TH-Gal4* expression^8,9^. These PAM-DAN are required for appetitive olfactory conditioning. We therefore wanted to know whether these PAM-DAN contributed to alcohol aversion, as indicated by our above pharmacological results, or whether they were involved in alcohol preference as suggested by their requirement for appetitive olfactory learning. When we activated the PAM-DAN subpopulation, using the heat-sensitive cation channel TrpA, flies showed enhanced experience-dependent alcohol-preference (Fig. 1f). Conversely, silencing these PAM-DAN using a dominant-negative dunamin, *shi*^*ts*^, abolished alcohol preference (Fig. 1f). These data suggested that *TH-Gal4* DAN mediate alcohol aversion, while PAM dopaminergic activity is required for experience-dependent alcohol preference. We confirmed the requirement for dopaminergic activity by knocking down the TH enzyme specifically in PAM-DAN (using *TH-RNAi*), which also abolished alcohol preference upon pre-exposure (Fig. 2b). This result also suggested that it is dopamine in these PAM neurons that is involved in experience-dependent alcohol preference rather than other putative co-transmitters.

**Figure 2:**
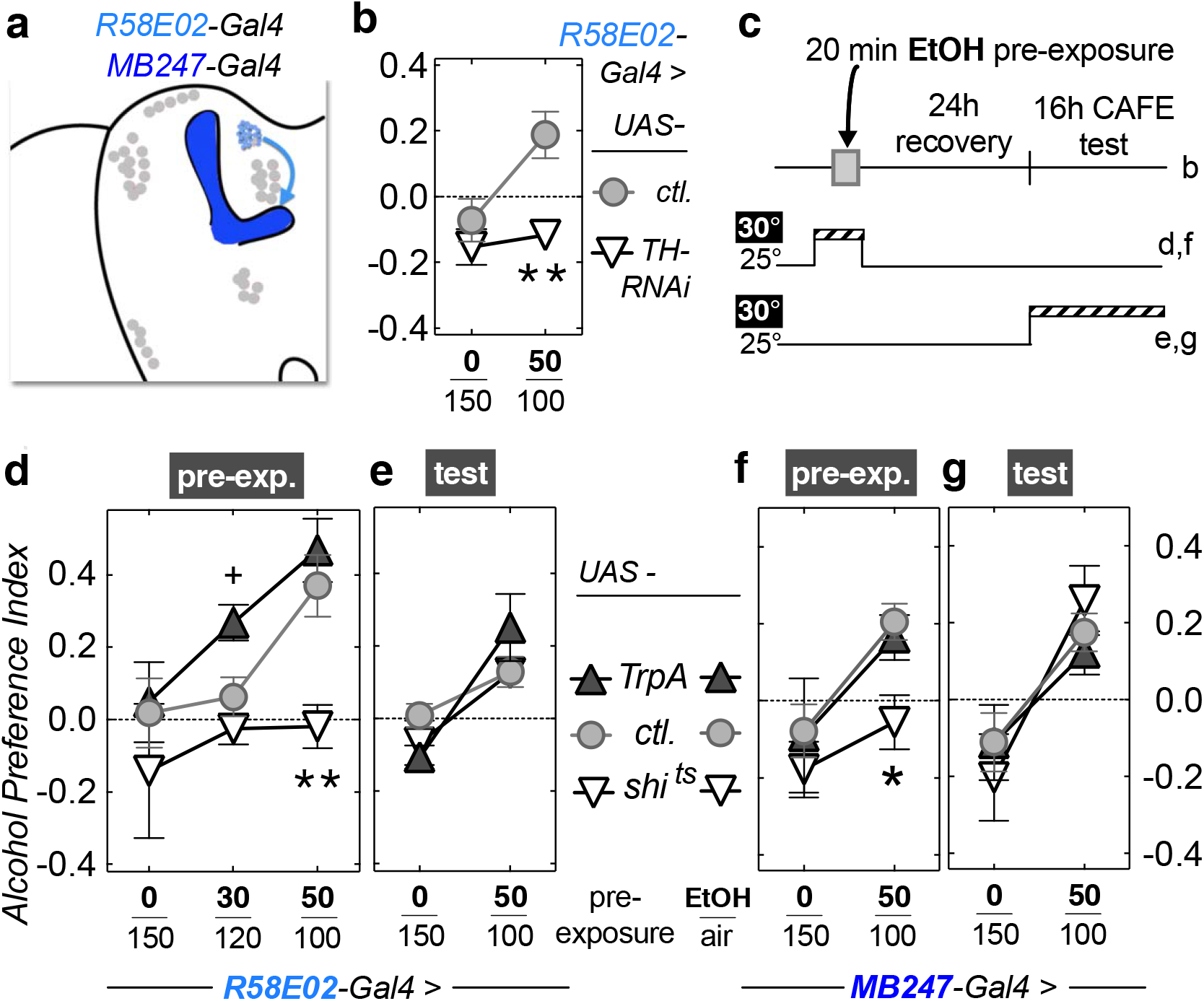
PAM dopamine neurons innervating the mushroom bodies are required for the acquisition of experience-dependent alcohol preference. (**a**) Fly brain schematic indicating the two drivers used. The mushroom body is indicated by dark blue. (**b**) RNAi-mediated knockdown of TH (*UAS-TH-RNAi*) in *PAM-Gal4* dopamine neurons (white) prevents experience-dependent alcohol preference (\*\**P* < 0.01). (**c**) Experimental paradigm used in d-g. (**d,e**) Silencing *PAM-Gal4* dopamine neurons with *UAS*-*shi*^*ts*^ during pre-exposure (d, white), but not during testing (e) leads to loss of experience-dependent ethanol preference (\*\**P* < 0.01). Activating PAM neurons during pre-exposure with *UAS-TrpA* (d, black) also shows a trend for facilitation of preference acquisition (^+^*P* = 0.056). (**f**,**g**) Similar to PAM neurons, silencing the target mushroom body neurons (*MB247-Gal4* driver) with *UAS-shi^ts^* during pre-exposure (f, white), but not during testing (g) also prevented formation of experience-dependent alcohol preference (\**P* < 0.05).

### PAM dopamine neurons are required for the acquisition of experience-dependent alcohol preference

Because our behavioral paradigm consists of an alcohol pre-exposure followed by a test of flies’ consumption preference a day later, we were able to ask whether PAM-DAN are required during the acquisition or the expression of alcohol preference. Limiting the heat-induced activation and silencing of the PAM-DAN to the alcohol pre-exposure or to the CAFÉ test revealed that these neurons had no effect during the testing (Fig. 2e). However, PAM-DAN activity was required during pre-exposure to acquire alcohol preference (Fig. 2d), and activating those neurons showed a trend towards facilitated acquisition of preference (Fig. 2e, *P* = 0.056). PAM-DAN project to the mushroom bodies (MB) a known center for associative learning in flies (Fig. 2a^8,9^). We therefore asked whether MB neurons are involved in experience-dependent alcohol preference, and, if so, when? Inhibiting activity of the α/β and γ lobes of the MB during the pre-exposure abolished preference acquisition (Fig. 2f), while we found no effect of the MB α/β and γ lobe neurons during the CAFÉ preference test (Fig. 2g). MB-projecting PAM-DAN activity is therefore required for the acquisition of alcohol preference, similar to the nucleus accumbens-projecting mammalian ventral tegmental area DAN^2^.

### A PPL1-to-fan-shaped body circuit mediates naïve alcohol aversion

We next investigated the role of the *TH-Gal4* DAN population (Fig. 3a) in naïve alcohol aversion. Knocking down *TH* expression in these neurons resulted in naïve alcohol preference (Fig. 3b), suggesting that dopamine is the relevant neurotransmitter mediating naïve alcohol aversion. Activation or silencing of the *TH-Gal4* DAN during ethanol pre-exposure had no effect on alcohol preference (Fig. 3c), but activating them during the CAFÉ preference test enhanced naïve aversion and abolished experience-dependent preference (Fig. 3d). Together with the data from Fig. 1b, e, these results suggested that alcohol acutely induces aversion via *TH-Gal4* DAN during the CAFÉ preference choice. This could be mediated by alcohol odor and/or taste sensation, but in addition, *TH-Gal4* DAN might also be directly activated by alcohol’s acute direct effect on these DAN.

**Figure 3:**
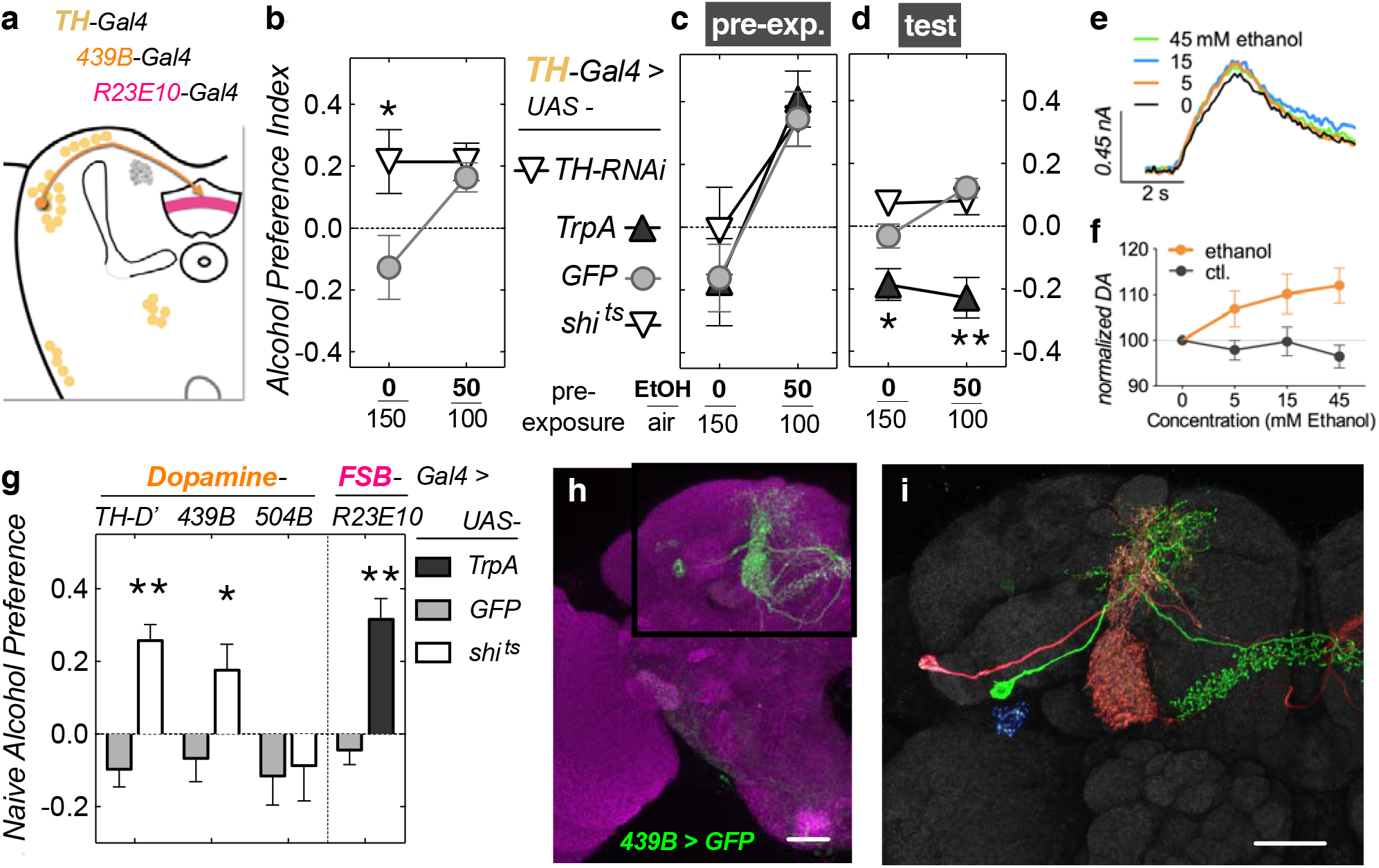
PPL1 dopamine neurons projecting to the fan-shaped body mediate acute naïve alcohol avoidance. (**a**) Schematic highlighting dopamine cells (*TH-*, *439B-Gal4*) and their target, the fan-shaped body (*R23E10-Gal4*). (**b**) Knockdown of TH (*UAS-TH-RNAi*, white) in *TH-Gal4* neurons leads to naïve preference (\**P* < 0.05). (**c**) Activating or silencing *TH-Gal4* dopamine neurons during ethanol pre-exposure does not affect preference, while activating them during the test (**d**) increases alcohol avoidance, regardless of pre-exposure. (**e**) Representative voltammetry traces from a larval brain and (**f**) total dopamine released upon stimulation of *TH-Gal4* neurons (with red light, via *UAS-CsChrimson*) shows that even low levels (5 mM) of acute ethanol potentiate dopamine release. Dopamine was measured from the median protocerebrum, close to the fan-shaped body. (**g**) A subset of *TH-Gal4* dopamine neurons affects naïve alcohol avoidance when silenced during naïve preference testing (\*\**P* < 0.01, \**P* < 0.05), including *TH*-*D’-Gal4*, which expressed in the PPL1 cluster, and *439B-Gal4*, which expresses in three PPL1 dopamine neurons (**h**). (**i**) Higher magnification picture of two (pseudocolored) *439B-Gal4* neurons. Two dopamine neurons PPL1-γ2α’1 (red) and PPL1-α’2α2 (cell body not visible in this picture) project to the mushroom bodies, and are also contained in the *504B-Gal4* driver, which does not affect naïve alcohol avoidance (see **g**). The green dopamine neuron (PPL1–dFSB) projects to the dorsal FSB, but not the MB. Activating layer 6 dFSB neurons (*R23E10-Gal4* driver) leads to naïve alcohol preference (see **g**).

We tested this latter hypothesis by performing *ex vivo* dopamine voltammetry experiments, while using the red light-activated CsChrimson cation channel^10^. Light-induced activation of *TH-Gal4* DAN caused robust release of dopamine from larval brain explants (Fig. 3e). Exposure to increasing amounts of alcohol revealed that even a low alcohol concentration of 5 mM potentiated the release of dopamine (Fig. 3e,f). This level is less than one third of the legal blood alcohol limit for driving. Acute direct exposure of *TH-Gal4* DAN to alcohol may therefore contribute to their activation, and underlie or potentiate acute sensory-induced aversion to alcohol at the behavioral level. Similarly, PAM-DAN might also be activated at these doses and thereby cause an appetitive reinforcing signal to the MB, ‘teaching the flies’ to like alcohol and override its aversive effects. This model would also explain how a sub-threshold ethanol pre-exposure could induce preference when paired with TrpA-mediated activation of PAM-DAN (Fig.2d, 30/120 EtOH/air flow; below threshold to turn aversion into preference in control flies).

One of the *TH-Gal4* DAN clusters activated by aversive stimuli is PPL1 (protocerebral posterior lateral 1)^6^, and these DAN might therefore also be involved in alcohol aversion. We thus followed up on other Gal4 drivers expressing in PPL1 to ask which of the *TH-Gal4* DAN are important in naïve, acute alcohol aversion. Using a number of Gal4 lines expressed in smaller subsets of *TH-Gal4* and PPL1 DAN (Supplementary Fig. 2)^11,12^, we found that silencing of both *TH-D’-Gal4* and *439B-Gal4* DAN caused naïve alcohol preference (Fig. 3g). *439B-Gal4* is expressed in 3 DAN per hemisphere (from the PPL1 DAN cluster, Fig. 3h), two projecting to the MB (Fig. 3i, red example), and one projecting to the dorsal fan-shaped body (dFSB; Fig. 3i, green). The two MB-projecting 439B-DAN are also expressed in line *504B-Gal4*^12^, which had no effect on acute alcohol aversion (Fig. 3g). This suggests that the dFSB-projecting 439B-DAN is involved in acute alcohol aversion. We further tested the involvement of dFSB in alcohol aversion by activating the PPL1-dFSB target neurons^13^ and found that this also caused naïve alcohol preference (Fig. 3g). Interestingly, this PPL1-dFSB circuit is involved in arousal from sleep^11,13^. Note that flies with silenced PPL1-DAN, or activated dFSB neurons consumed the same total amount of food as did controls (Supplementary Fig. 3), arguing that these manipulations did not cause pervasive ‘sleepiness’ or inactivity (Supplementary Fig. 4).

### A Dopamine D1-like receptor is required in dopamine target neurons for alcohol aversion and preference

Together, the above data suggested that the PPL1-dFSB circuit mediates acute alcohol aversion, while the PAM-MB circuit participates in the acquisition of experience-dependent alcohol preference. To further strengthen our evidence for a critical role of dopamine in both these circuits, we investigated the involvement of two major dopamine receptors, D1 and D2. Pan-neuronal knock down (Fig. 4a) of the D1-like Dop1R1 (*D1R*) receptor led to naïve alcohol preference (Fig. 4d), while knock down of the D2-like receptor (*D2R*) did not affect naïve aversion or experience-dependent preference (Fig. 4e). Furthermore, knock down of *D1R* in the dFSB was sufficient to cause naïve alcohol preference (Fig. 4g). MB-specific knock down revealed that *D1R* is required in MBs for experience-dependent preference (Fig. 4f).

**Figure 4.**
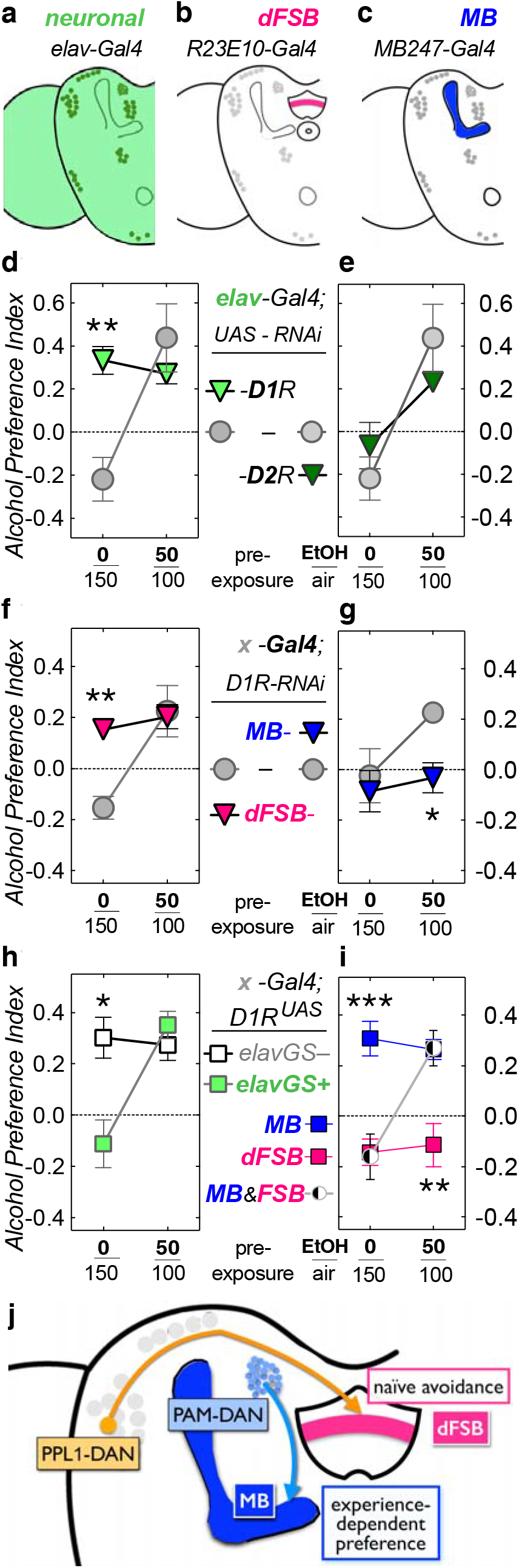
The dopamine D1R1 receptor is required in the mushroom bodies for experience-dependent alcohol preference and in the fan-shaped body for naïve avoidance. (**a**-**c**) Fly brain schematics showing Gal-4 drivers used expressing in all neurons (**a**), dorsal fan-shaped body layer 6 (dFSB, **b**), and mushroom bodies (MB, **c**). (**d**) Pan-neuronal, RNAi-mediated knockdown of dopamine D1R1 receptor (*D1R*), but not D2 receptor (*D2R*, **e**) abolishes naïve alcohol avoidance (\*\**P* < 0.01). (**f,g**) D1R knockdown in the dFSB also abolishes naïve alcohol avoidance (\*\**P* < 0.01, magenta in **f**), whereas, D1R knockdown in the MB abolishes experience-dependent preference (\*\**P* < 0.05, blue in **g**). (**h,i**) *D1R*^*UAS*^ flies are mutants that lack *D1R* expression, but this can be restored by the introduction of a Gal4-driver. Together with an RU486-inducible pan-neuronal Gal4 driver (*elavGS*, for RU486-Gene-Switch), these *D1R*^*UAS*^ mutants show naïve alcohol preference when Gal4 is not induced (*elavGS–*, white in **h**). This is rescued when adult flies are fed RU486 before the ethanol-pre-exposure and D1R expression is restored (*elavGS+*, green in **h**). When *D1R* expression is restored in the MB, *D1R*^*UAS*^ flies still show naïve alcohol preference (*MB247-Gal4>D1R*^*UAS*^, blue in **i**). Conversely, restoring *D1R* expression in the FSB only rescues naïve alcohol avoidance, but now reveals a loss of experience-dependent preference (*R23E10-Gal4>D1R*^*UAS*^, magenta in **i**), presumably due to lack of *D1R* in the MB. Indeed, when *D1R* expression is restored in both the MB and the FSB, *D1R*^*UAS*^ mutants show normal naïve alcohol avoidance succeeded by preference upon an alcohol pre-exposure (*MB247-Gal4;R23E10-Gal4>D1R*^*UAS*^, black and white in **i**). (**j**) Schematic highlighting the two dopamine circuits involved in naïve alcohol avoidance (PPL1–dFSB) and the acquisition of experience-dependent alcohol preference (PAM–MB).

To ascertain that our results were not caused by RNAi off-target effects, we also tested the *D1R*^*UAS*^ mutation. These flies lack functional *D1R* expression, but the presence of a Gal4-binding UAS site in the *D1R* gene allows expression to be restored by a Gal4 driver ^14^. As with pan-neuronal RNAi knock down of *D1R* (Fig. 4d), *D1R*^*UAS*^ flies showed naïve alcohol preference (Fig. 4h). When we conditionally restored adult *D1R* expression in neurons using an inducible pan-neuronal Gal4-driver (*elavGS*+), flies showed normal naïve alcohol aversion, which turned into experience-dependent preference upon pre-exposure (Fig. 4h). These data provide evidence for an acute requirement of dopamine signaling during adult behavior (as suggested by Figs. 2, 3) and argue against developmental defects causing the behavioral deficits^15^. When we restored MB-specific *D1R* expression in *D1R*^*UAS*^ flies, we observed no change from the *D1R*^*UAS*^ mutants, i.e., naïve preference with intact experience-dependent preference (Fig. 4i). Conversely, dFSB-specific *D1R* expression rescued *D1R*^*UAS*^ naïve preference back to normal naïve avoidance, but these dFSB-rescued *D1R*^*UAS*^ mutants did not acquire experience-dependent preference (Fig. 4i), presumably due to the lack of D1R in the MB. Indeed, when we restored D1R expression in both the MB and dFSB, *D1R*^*UAS*^ flies displayed normal naïve alcohol aversion, followed by experience dependent preference upon alcohol pre-exposure (Fig. 4i). These data confirm that dopaminergic signaling from the PAM to the MB is required for acquired alcohol preference, and PPL1 dopaminergic signaling to the dFSB is required for naïve alcohol avoidance (Fig. 4j).

## Discussion

Here we demonstrate the multifaceted involvement of dopamine in voluntary alcohol consumption. We show that PAM dopamine neurons are required for the acquisition of experience-dependent alcohol preference. Consummatory preference also requires the dopamine D1R1 receptor in the *Drosophila* MB. This PAM to MB circuit has previously been found to be important in the acquisition of appetitive sucrose learning^8,9^, while D1R1 is well known to be involved in *Drosophila* learning and memory^16^. Furthermore, the MB are involved in ethanol-reinforced odor preference^4^, as well as preferential alcohol consumption^17^. Our findings that acute alcohol exposure potentiates dopaminergic release suggest a mechanism of how an acute vapor exposure of ethanol might cause reinforcement: as alcohol rises in the brain, dopaminergic neurons, including reinforcing PAM neurons, get activated and impart an association of cues with behavioral reinforcement. These cues might involve the smell of alcohol itself, but this remains to be determined. Still, our data suggest that the Drosophila PAM neurons act analogously to the ‘classical’ mammalian dopamine neurons projecting from the ventral tegmental area to the nucleus accumbens, thought to be important in mediating the reinforcing actions of drug of abuse.

Our data also suggests that a single bilateral pair of PPL1 dopamine neurons mediates acute aversive effects of ethanol, and silencing these neurons causes abolished alcohol aversion. Dopaminergic neurons in the PPL1 cluster are acutely activated by a number of aversive stimuli^6^, consistent with their involvement in alcohol aversion. Thus in our acute choice paradigm, the smell^18^, or taste^19,20^ of ethanol might activate these neurons and contribute to acute avoidance of alcohol consumption. In addition, acute drinking and alcohol’s pharmacodynamic potentiation of these neurons might also contribute to acute avoidance. Interestingly, the PPL1-dFSB circuit is involved in arousal (from sleep)^11,13^, and our findings suggest that this circuit might be more broadly involved in the processing and salience of aversive cues.

Taken together, our data show that *Drosophila* dopamine neurons are required for both reinforcing and acute aversive reactions to alcohol. This is similar to recent findings in the mammalian brain, where dopaminergic neurons are involved in opposing reactions to drugs of abuse^21,22^. We show that alterations in either of these dopamine circuits can lead to a reduction in ethanol intake, emphasizing that acute sensitivity to the aversive aspects of a drug are protective against the development of addiction. This is in line with human genetic findings, where the strongest genetic factors that protect people from developing alcohol abuse disorders are alcohol dehydrogenase variants, which are involved in triggering acutely unpleasant reactions^23^. Furthermore, people perceiving alcohol as acutely more bitter tasting, are also less likely to develop alcoholism^24,25^. *Drosophila* thus show complex reactions to ethanol that are similar to humans. In addition, distinct *Drosophila* dopaminergic circuits mediate these diverse reactions. Thus, while the structural architecture of the fly brain is clearly different from that of mammals, key sub-circuits are conserved in their logic and in their multifaceted use of the same neurotransmitter—dopamine—which can mediate opposing behavioral outcomes, depending on the specific dopaminergic circuit engaged.

## Materials and Methods

### Fly husbandry and Genetics

*Drosophila melanogaster* were raised in a 12:12 hr. Light/Dark cycle on a standard cornmeal/molasses diet at 25°C with 70% humidity, except for temperature sensitive experiments, which used 30°C during the experiments. The genetic background for all experiments used was *white* Berlin* (unless explicitly stated). Transgenic Gal4 driver lines containing different regions of the TH genomic locus (*TH-C’-, D’-, D4-, F2-*, and *C1-Gal4*) were obtained from Dr. Mark Wu (John’s Hopkins). PPL1 specific-Gal4 lines were obtained from Dr. Karla Kaun (Brown University). Other transgenic lines were obtained from the Bloomington stock center.

### Drug feeding protocols

Pharmacological treatment with 3-iodo tyrosine (3IY, Sigma) and L-DOPA (Sigma) were carried out as previously described^26^. 3IY (10mg/ml) or L-Dopa (1mg/ml) were dissolved in solutions containing 250 mM sucrose. Flies were pre-fed in a modified CAFÉ assay in rectangular 4-well plates (128 × 85.5 mm, Thermo Scientific; Fig. 5). Food was provided in 0.2 ml tubes with a 27 G needle hole at the bottom for drinking access, a 27 G hole atop for pressure equilibration and a 25 G hole atop for filling with solution. Flies were fed 3IY for a period of 48-hours and L-DOPA for 24-hours in the modified CAFÉ apparatus. For the *elav-GeneSwitch* Gal4 experiments, food-deprived flies were fed with 0.5 mM mifepristone (RU486) for 3 hours prior to pretreatment to ethanol.

### Booz-o-mat exposure

Exposure paradigms used are as previously described^3^. The day before ethanol vapor exposure, male flies were collected in groups of 30 and put on un-yeasted food. The following day, flies were transferred into the Booz-o-mat apparatus for a 20-minute exposure at desired ethanol to air ratio (E/A) as described^27^. For temperature-sensitive experiments, *UAS*-*shi^ts^, UAS*-*TrpA^ts^*, and control flies were allowed to acclimate at 30°C for 20 min in the Booze-o-mat before starting the 20-min exposure at 30°C. Flies were then transferred to vials and placed into a 25°C/70% humidity incubator for a 24-hour recovery period.

### Capillary Feeder (Café) assay

24 hr after ethanol pre-exposure, 15 flies were placed into each well of the Café assay apparatus as described^28^. Preference testing was carried out for 16-hours.

### Immunohistochemistry

Immunohistochemistry was performed as previously described^29^. To visualize *439B-PPL1-GAL4* expression in the brain, a *pJFRC225-5×UAS-IVS-myr::smGFP-FLAG* (*smGFP-FLAG*) reporter probe^30^ was utilized. The smGFP-FLAG transgene was visualized with an anti-FLAG antibody. The presynaptic marker mouse anti-nc82 antibody was used to label general neuropil/brain structure. The multicolor Flip-out approach^31^ was used for stochastic labeling of 439B-PPL1 neurons.

### Light-induced stimulation of DA neurons and fast-scan voltammetry

All chemicals were purchased from Sigma-Aldrich (St. Louis, MO) and all solutions were made with Milli-Q water (Millipore, Billerica, MA). Dissections, recordings, and calibrations were performed in a simple buffer solution (131.3 mM NaCl, 3.0 mM KCl, 10 mM NaH_2_PO_4_ monohydrate, 1.2 mM MgCl_2_ hexahydrate, 2.0 mM Na_2_SO_4_ anhydrous, 1.2 CaCl_2_ dihydrate, 11.1 mM glucose, 5.3 mM trehalose, pH = 7.4). Carbon fiber microelectrodes were fabricated from T-650 carbon fibers (a gift of Cytec Engineering Materials, West Patterson, NJ) and were used for fast-scan cyclic voltammetry as previously described^32^. Virgin females with *UAS-CsChrimson* (a chimera of CsChR and Chrimson) inserted in attp18^33^ (a gift of Vivek Jayaraman) were crossed with *TH-GAL4* (a gift of Jay Hirsh). Resulting heterozygous larvae were shielded from light and raised on standard cornmeal food mixed 250:1 with 100 mM all-trans-retinal. A small amount of moistened Red Star yeast (Red Star, Milwaukie, WI) was placed on top of the food to promote egg laying.

For the protocerebrum recordings, brains were isolated into dissection buffer from larvae using forceps under a dissection microscope, and the electrode was implanted from the lateral edge of the tissue into the dorsal medial protocerebrum. The electrode equilibrated in the tissue for 10 minutes prior to data collection and a baseline recording was taken for 10 seconds prior to stimulation. Red light estimated at 0.75 mW from a 617 nm fiber-coupled high-power LED with a 200 μm core optical cable (ThorLabs, Newton, NJ) was used to stimulate the CsChrimson ion channel. The TarHeel CV software (a gift of Mark Wightman) was used to control the light stimulation and to record the current from the applied voltage. After taking a baseline 2 second stimulation, 5 mM ethanol (10% in buffer) was added to the solution of fly buffer and then another stimulation was recorded after 5 minutes. The concentration of ethanol was increased to 15 mM and then to 45 mM. Stimulations were performed at each concentration five minutes after the ethanol was added to allow for equilibration. Adding increasing amounts of dissection buffer instead of ethanol was performed as a vehicle control. Data are presented at mean +/− standard error of the mean (SEM) and graph error bars are SEM.

### HPLC measurement of brain dopamine levels

Flies were immobilized on ice (5/sample). Brains were dissected and homogenized in ice-cold 0.02 M HClO_4_ and 0.00025 M ascorbate solution using a Kontes micro tissue grinder. Homogenate was centrifuged at 13,000 rpm for 5 min at 4ºC. The supernatant was collected and centrifuged at 13,000 rpm for 30 sec at 4°C using a 0.22 µm Ultrafree PVDF filter (MilliporeSigma, Billerica, MA). Samples (approx. 25 µL final volume) were stored at −80ºC until analysis. The HPLC-ECD system consisted of a Luna 3 µm C18 (2) 100 Å, 50 × 1 mm, LC column (Phenomenex, Torrance, CA) and SenCell2 electrochemical cell at +450 mV (Antec Leyden, Netherlands) with Ag/AgCl reference electrode. Aqueous mobile phase with ion-pairing agents (0.50 g OSA, 0.05 g DSA, 0.13 g EDTA, 11.08 g NaH_2_PO4, 100-150 mL MeOH in 1L H_2_O; pH adjusted to 5.6) was delivered to the column using a LC110S piston pump (Antec Leyden, Netherlands). Column and electrochemical cell were kept at 35ºC inside a controller (Intro model, GBC Separations, Hubbardson, MA) with a Rheodyne 8125 manual injector. Analog responses from the electrochemical detector were digitized using an analog-to-digital converter (SS240× model, Scientific Software, Pleasanton, CA). EZ Chrome Elite software was used to collect chromatograms (Scientific Software, Pleasanton, CA). Sample dopamine content was estimated via the calibration curve method using external dopamine standards prepared in 0.1 M H_3_PO_4_. Reported estimates are derived from chromatograms with signal-to-noise ratio ≥ 3 that met in-house quality criteria.

### Statistics

Statistical significance of results in this manuscript was established using analyses of variance (ANOVAs) tests with GraphPad Prism software for Mac. For the post-hoc analyses, Dunnett’s test was applied to control for the multiple comparison when several groups were compared to the same control. Error bars in all experiments represent SEM. Significance was only attributed to experimental lines that were statistically different from their respective controls. Significance in all graphs shown are defined as *p < 0.05, **p < 0.01, and ***p < 0.001.

## Acknowledgements

We thank Dante Gonzalez, Rachel Dove, Summer Acevedo and the rest of the Rothenfluh lab for their help and advice with experiments. We thank Mark Wu for numerous dopaminergic *Gal4* flies and Karla Kaun for the PPL1-Specific Gal4 lines. This work was supported by the NIH (T32 fellowship DA007290 and F31 AA021340 to SAO, T32 fellowship AA007471 to RUC, R37AA011852 to RAG, R01DK110358 to ARR and R01AA019526 to AR). AR was also supported by an Effie Marie Cain Scholarship in Biomedical Research from UT Southwestern, and by the University of Utah Neuroscience Initiative. ARR was also supported by the AHA (16CSA28530002). The authors declare that they have no competing interests. A detailed description of all material and methods as well as supplementary figures are available as supporting online materials.

## Author Contributions

S.A.O. and A.R. conceived the study. S.A.O., E.P.C., Y.N., D.W., R.A.G performed experiments and analyzed the data with the other authors. R.A.G., G.M.R., B.J.V., A.R.R., A.R procured funding. S.A.O., A.R.B., C.B.M., A.R.R., and A.R. wrote the paper with input from the other authors.

---

**Supplemental Figure 1.**
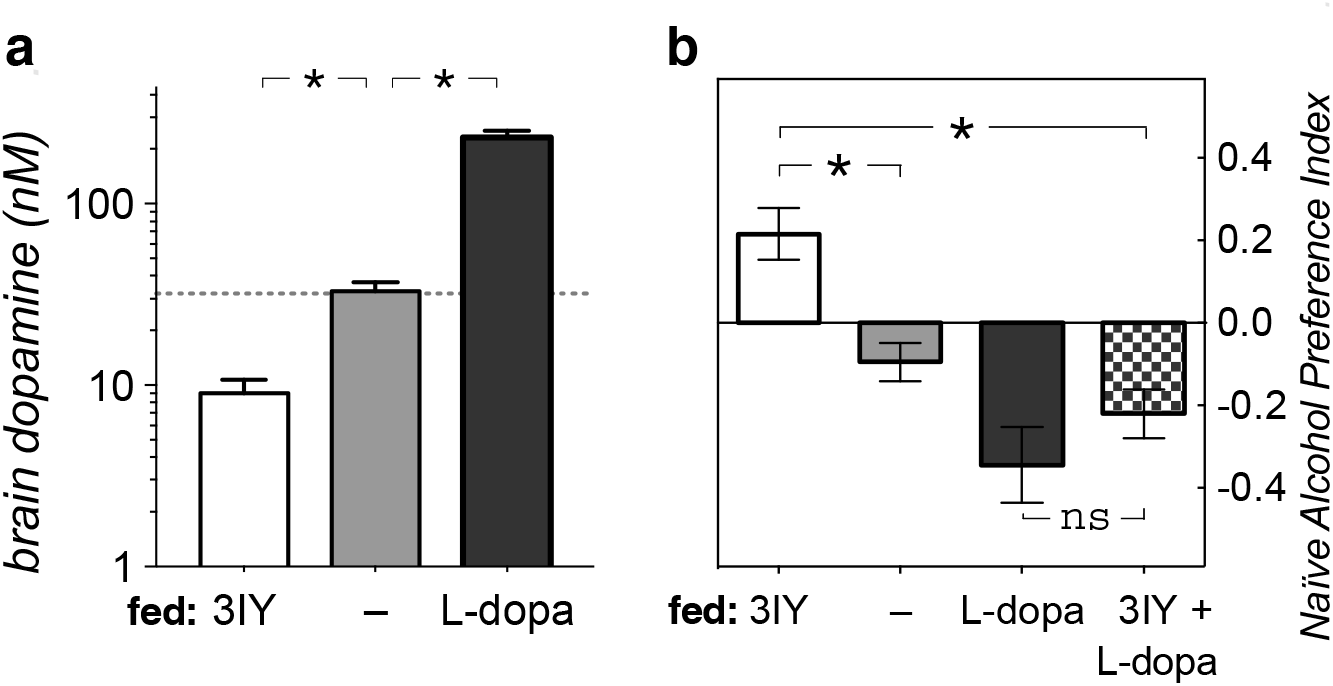
Dopamine pharmacology. (**a**) Brain dopamine concentrations in pooled fly brain homogenates after drug feeding, as measured by HPLC-ECD (*\*P* < 0.05, n = 3 replicates of 5 brains homogenized). (**b**) Simultaneous feeding of 3IY and L-dopa converts 3IY-induced naïve preference back to naïve avoidance. (Data as in main Fig. 1B, with the addition of the co-feeding, checkered; *\*P* < 0.05 one-way ANOVA with Dunnett’s post-hoc comparison).

**Supplementay Figure 2.**
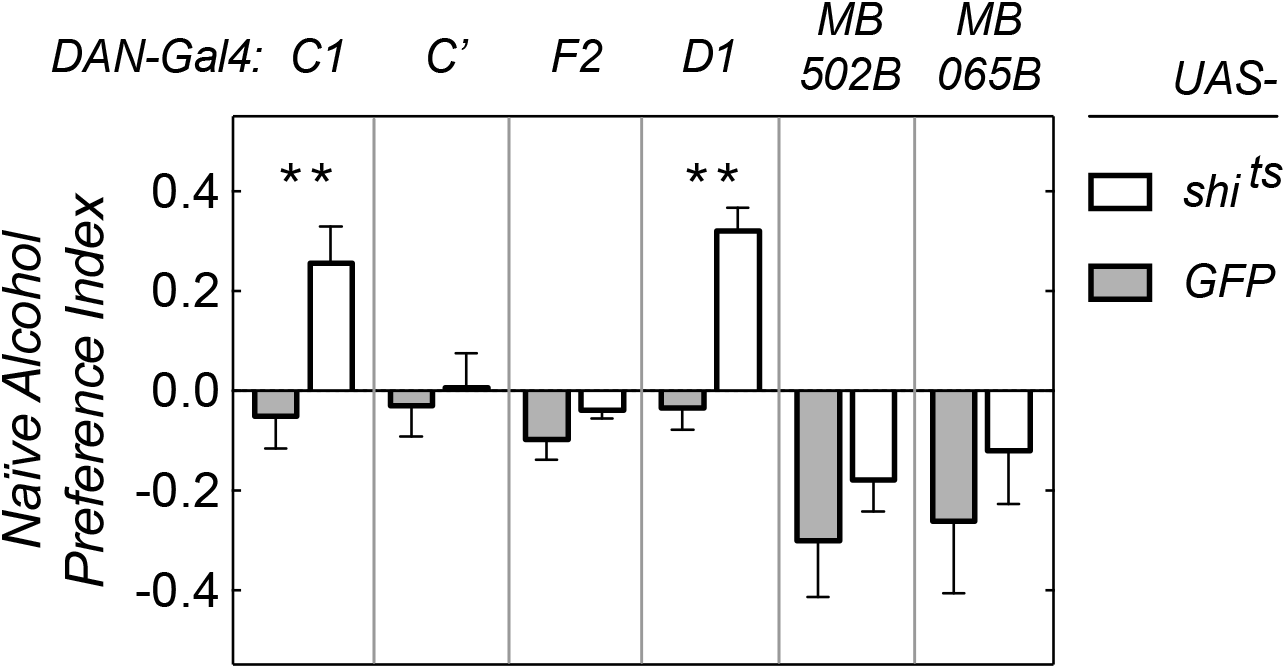
Silencing of different DAN causes naïve alcohol aversion. 2 of 6 transgenic *DAN-Gal4* driver lines (in addition to the ones shown in Fig. 3g) caused naïve preference to alcohol when silenced with *UAS-shi^ts^* compared to their respective *UAS-GFP* controls (*\*P* < 0.05, one-way ANOVA with Bonferroni correction, n = 6–12).

**Supplementay Figure 3.**
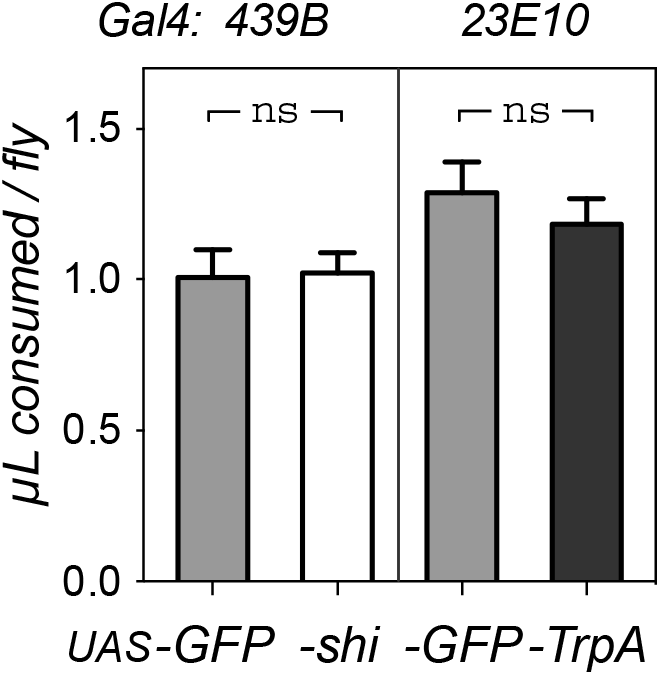
Naïve alcohol preference caused by manipulation of the PPL1– dFSB circuit does not alter total food consumption. Total food consumption (sucrose + sucrose/ethanol mixture) per fly is indicated over the 16-hour CAFÉ test. No significant differences were observed between experimentals and controls (grey). This was true for other DAN-Gal4 lines affecting naïve alcohol aversion too (data not shown).

**Supplementay Figure 4.**
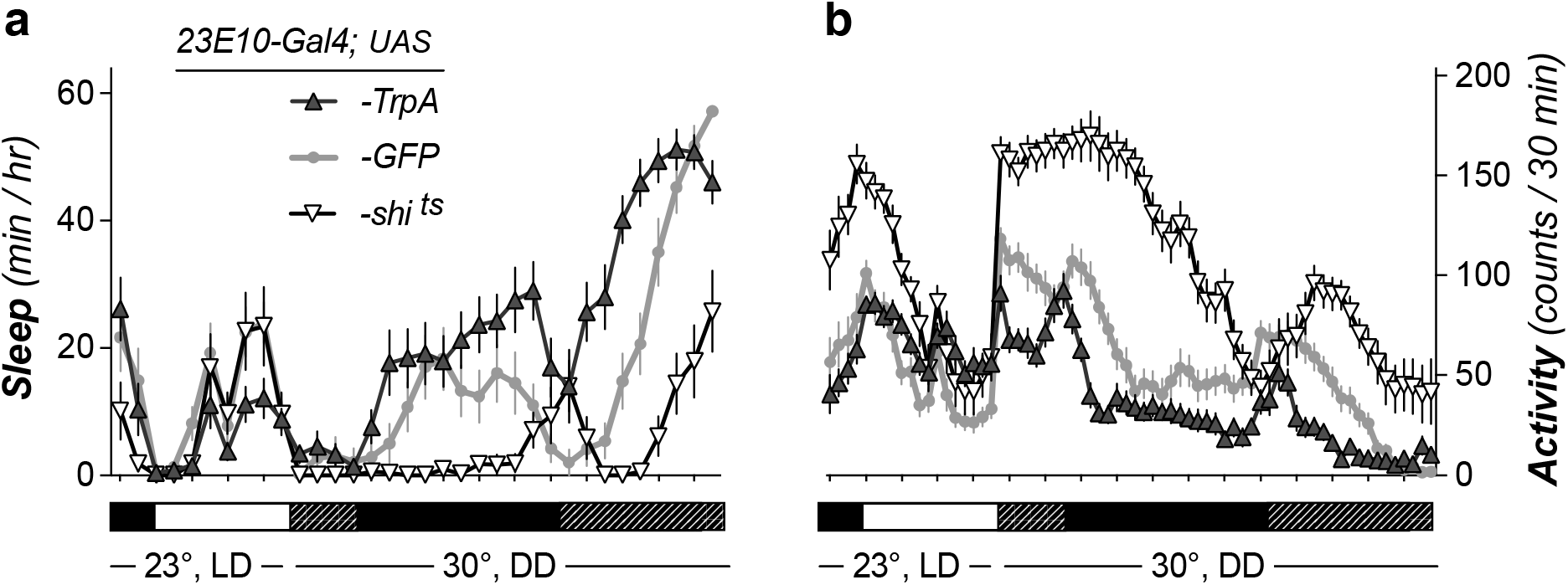
Sleep induction by activation of dFSB. (**a**) Shifting *23E10-Gal4; UAS-TrpA* to 30°C (black) caused an increase in total sleep duration, but did not just put the flies to sleep: considerable levels of activity remained (**b**). Note that this is the same temperature and light regimen as used for the CAFÉ preference test.

